# Optical Phenotyping Using Label-Free Microscopy and Deep Learning

**DOI:** 10.1101/2025.04.01.646480

**Authors:** Shuyuan Guan, Thomas Knapp, Alba Alfonso-Garcia, Suzann Duan, Travis W. Sawyer

## Abstract

**Significance:** Tissue phenotyping plays a critical role in biomedical research and clinical applications by providing insight into the structural and functional characteristics of tissues that can characterize clinical behavior and identify therapeutic targets. However, conventional phenotyping techniques are destructive, time-intensive, and expensive, posing challenges for both efficiency and widespread use.

**Aim:** To develop an optical phenotyping approach in pancreatic cancer specimens using label-free multiphoton microscopy combined with spatial transcriptomics and deep learning.

**Approach:** We measure and co-register a dataset comprised of spatial transcriptomics, autofluorescence, and second harmonic generation microscopy. We then cluster tissue subregions into meaningful phenotypes using transcriptomic signatures. We then evaluate three different classification models to predict phenotype based on label-free imaging data, and assess generalizability and prediction accuracy.

**Result:** Our deep-learning classification model achieves over 89% accuracy in classifying six tissue types using label-free microscopy images. The one-vs-rest AUC values for all classes approaches 1, confirming the robustness of our model.

**Conclusion:** We demonstrate the feasibility of optical phenotyping in distinguishing the structural and functional characteristics of pancreatic cancer specimens. Integrating additional gene-expression data or complementary label-free imaging modalities, such as fluorescence lifetime imaging microscopy, holds the potential to further enhance its accuracy and expand its applications in clinical research and diagnostics.

**Statement of Discovery:** We develop and demonstrate a method for optical phenotyping using deep learning to classify label-free microscopy images into tissue phenotypes defined by transcriptomic signatures.

## 1 Introduction

Tissue phenotyping plays a decisive role in profiling disease progression and onset, particularly in the context of cancer.^1^ For example, many phenotyping efforts have focused on profiling the tumor microenvironment (TME), which plays an important role in disease progression and metastasis.^2^ With the recent advancements in precision medicine, tumor phenotyping is a crucial step to identifying specific therapeutic targets for diseases that are potentially heterogeneous, for example in pancreatic cancer.^3^ Towards this end, phenotyping of ex vivo tissues, including those based on sequencing approaches, can increase our understanding of TME features that determine disease prognosis.^4^ While this provides incredible power for mapping tissue and cell genotype-phenotype relationships, an outstanding challenge is the lack of approaches that are compatible for in vivo tissue phenotyping. In addition, many such methods require significant processing time and can be costly - the ability to identify phenotypes in a minimally invasively fashion, and without lengthy and expensive processes, could offer opportunities to glean insight into fundamental processes of disease onset and evolution in basic science research, as well as could lead to unique approaches for advancing precision medicine. In this manuscript, we primarily focus on tissue phenotypes that are defined by gene expression as measured through RNA sequencing, or transcriptomics, as this provides a description of underlying biological processes that dictate tissue function.^5^

Label-free biomedical imaging techniques could provide an avenue for rapid, scalable, and/or in vivo compatible “optical phenotyping” by sensing naturally occurring contrast to visualize fundamental tissue properties, which provide powerful insight into biological processes and pathological events.^6^ For example, autofluorescence and fluorescence lifetime imaging can probe endogenous markers of metabolism, immune response, and vasculature.^7–10^ Other methods, such as polarized light imaging, second harmonic generation, or optical coherence tomography can visualize microstructural features such as collagen distribution and the extracellular matrix.^11–13^ Different disease phenotypes can exhibit clear alterations in these characteristics – for example, immunologically cold and hot tumors within a specific type of cancer respond differently to immunotherapies that rely on the presence of tumor immune infiltrate.^14^ Other examples include extracellular matrix alterations that promote tumor cell invasion and migration towards vasculature.^15^ As such, it follows that label-free imaging could be utilized for minimally invasive and rapid measurement of key tissue properties to monitor biological events such as disease onset, or to identify specific tissue phenotypes. However, while label-free imaging shows promise, the non-linear relationships between genetic pathway alterations that dictate phenotypes and the resulting downstream functional changes has inhibited the ability to accurately map the underlying mechanistic pathways related to label-free contrast. An added complexity is that tissue changes associated with different diseases and phenotypes can occur at multiple scales and are not represented fully by the abundance of a given molecule^16^ - the spatial organization and distribution of these molecules across different spatial scales carries significant biological information. As such, fully realizing the potential of label-free imaging for optical phenotyping requires the integration of more sophisticated approaches for image feature extraction.

Recent advancements in deep learning may offer a solution to mapping the non-linear sequence-function relationship,^17^ offering a unique opportunity to bridge the gap and perform optical phenotyping of tissues. Since the explosion of artificial intelligence (AI) in biomedical imaging, a myriad of research efforts have aimed to integrate AI methods into different applications across the field. One of the most common applications is for image classification, where AI models such as deep neural networks (DNNs) are trained to classify images with extremely high accuracy.^18^ DNNs offer the ability to extract multi-scale hierarchical spatial features in image data, offering exceedingly high performance for image analysis tasks. Binary, multi-class and multi-label classification problems have all been investigated with much success; however, one limitation to these approaches are that the classes and/or labels are generally established based on general tissue categories, often with a significant subjective nature (e.g. diseased, normal, or pathological grades).^19^ Diseases such as cancer present with significant heterogeneity within tumor types and disease grades, and this intratumor heterogeneity contributes to drastically different phenotypes and clinical outcomes.^20, 21^ These nuances are lost when grouped together qualitatively, severely limiting both the performance and utility of these approaches. Tissue phenotypes can be reliably established quantitively based on the transcriptomic signature, and developing technology to classify tissue into different phenotypes using label-free imaging would be a powerful tool in biomedical research, as it could enable rapid, nondestructive, minimally invasive, and high precision tissue assessment.^22^

In this work, we demonstrate the ability to perform optical phenotyping using label-free microscopy in a set of patient specimens of pancreatic cancer. We perform autofluorescence and SHG imaging through multiphoton microscopy (MPM) on these specimens, as well as spatial transcriptomics. We co-register the dataset so that transcriptomic signatures are aligned with a region of interest of the label-free imaging modalities. Using this co-registered dataset, we first cluster tissue regions into different tissue types and phenotypes, and define these by mapping gene expression to the Human Protein Atlas. We extract image features for each channel including abundance and Haralick texture features. We performed statistical analysis on label-free imaging contrast between clusters, and then built several classifiers to classify tissue using label-free images into phenotypes defined using transcriptomic signatures. Our results show that the abundance of different label-free imaging channels is not sufficient for differentiating tumor phenotypes. Image texture describing spatial distribution of abundance provides higher performance, and using unsupervised feature extraction coupled with DNN classification achieves over 90% accuracy in performing optical phenotyping. These results show the value of hierarchical multi-scale feature extraction for capturing relevant biological characteristics. These promising results lay the foundation for our future work that will focus on expanding our patient cohort, optimization of DNN architecture, investigating the generalizability to other organs and diseases, as well as validating these findings in fresh tissues.

## 2 Methods

### 2.1 Sample Preparation

This study was performed on four specimens collected from four patients who were diagnosed with grade 2 pancreatic neuroendocrine tumors (NETs) via pathological examination. The formalin-fixed paraffin-embedded (FFPE) specimens were obtained from the University of Arizona Tissue Acquisition and Cellular/Molecular Analysis Shared Resource, originally collected under IRB 0600000609. All specimens were tested for RNA quality by evaluating the RIN and DV200 score. All specimens yielded a DV200 score of at least 50, and a RIN score of at least 2.3.^23, 24^ The specimens were sectioned at a thickness of 10 microns onto standard microscope slides. Two serial sections for each specimen were used in this study: one was placed on a specific slide for spatial transcriptomics purchased from 10X genomics (Visium slide; shown on the left in Figure 1) and a second adjacent section was used for MPM.

**Fig 1.**
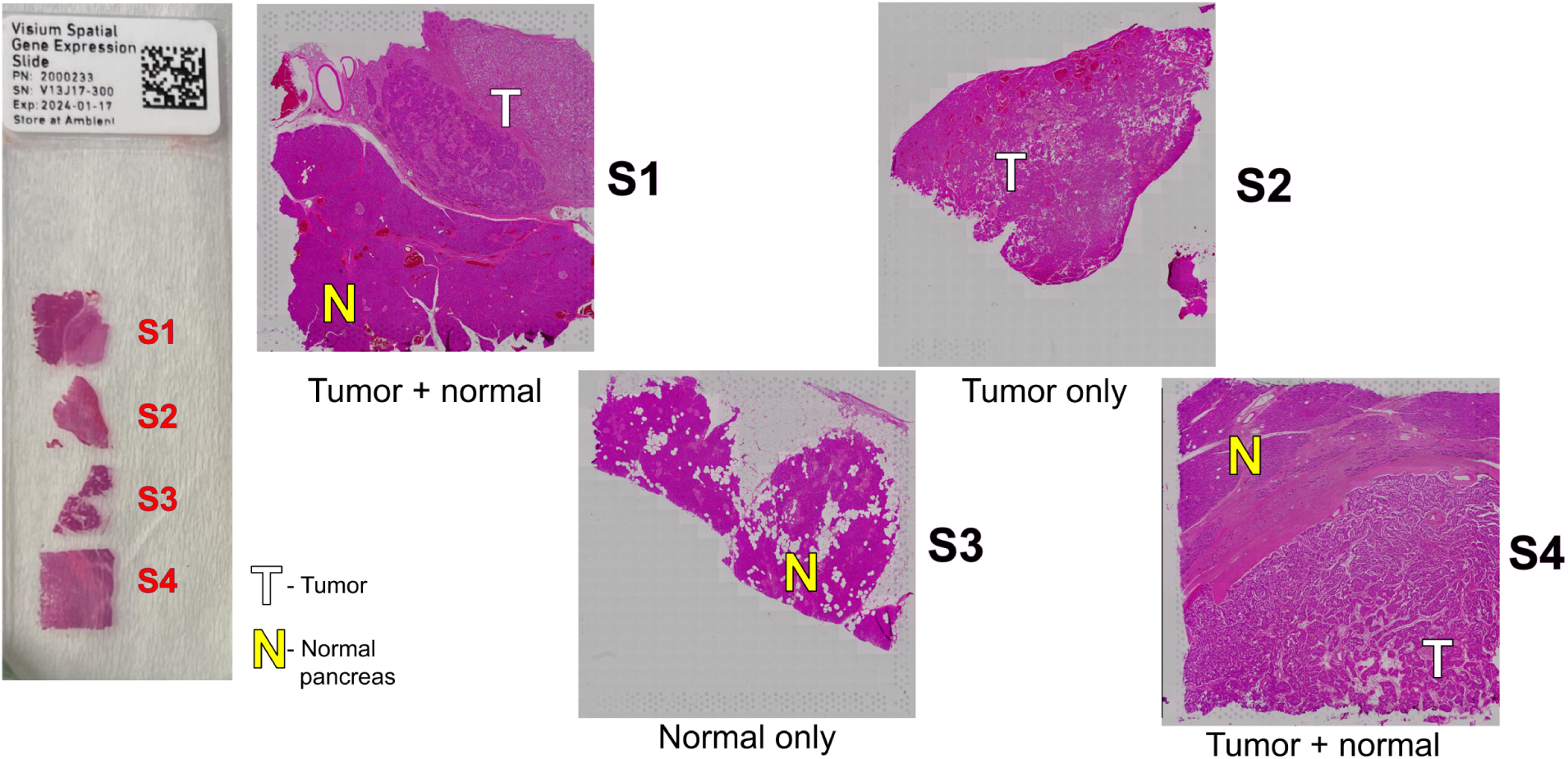
Diagram of pancreatic cancer patient specimens (S1-S4) used for the research study. Tumor and normal regions are denoted with T and N respectively. The left side of the figure shows the specimens placed on the Visium slide for spatial transcriptomics.

### 2.2 Imaging Data Collection

MPM imaging was performed on a single unstained histological section with a thickness of 7 microns using a Zeiss LSM 780 NLO system with an automated 3D translation stage. A tunable MaiTai Ti:Sa laser and GaAsP detectors were used to image in five spectral channels, four channels capturing the signal from tissue autofluorescence, and a second-harmonic generation (SHG) channel. The excitation and detection wavelengths are shown in Table 1 and have been used in previous studies.^25^ Each MPM channel is referred to in this manuscript by the endogenous molecule(s) believed to be the primary source of signal when imaging with the spectral parameters in Table 1. This terminology is meant to provide ease of reference, as multiple fluorescent species can provide overlapping spectra, and tissue fixation may alter endogenous markers. While these fluorescence properties could be elicited with single photon fluorescence, we use multiphoton to also elicit SHG simultaneously. 256 x 256 pixel tile scans were collected in a grid array over each sample and combined with post-processing to form a composite image of the full specimen as it was imaged on the Visium slide. A 20X objective was used, providing a square pixel resolution of 1.28 micron. Z-stacks were acquired for each tile scan to ensure in-focus signal was collected over the full range of the uneven tissue surfaces, with the number of z-stacks adjusted as necessary. Despite z-stacking, some uneven brightness within the individual tile scans results in a grid-like artifact in the image mosaics. Post-processing that has been previously described was used to balance image brightness between image tiles prior to quantification.^26^

**Table 1.** Imaging channels used for label-free microscopy. All images were acquired using two-photon excitation, thus the excitation wavelength is half of the listed wavelength.

| Channel | Two-photon Excitation Wavelength (nm) | Detection Wavelength band (nm) | Potential fluorophores / sources of contrast |
| --- | --- | --- | --- |
| 1 | 700 | 425-465 | NADH / Collagen |
| 2 | 920 | 475-600 | FAD |
| 3 | 830 | 550-600 | Lipofuscin / lipopigments |
| 4 | 800 | 590-625 | Porphyrin |
| 5 | 880 | 430-450 | Organized collagen (SHG) |

### 2.3 Spatial Sequencing

An adjacent histological section for each patient sample was cut onto the Visium slide before being deparaffinized and stained with Hematoxylin and Eosin (H&E) following the standard 10X Genomics protocol for FFPE tissue prior to being slide-scanned using a Nikon BioPipline SLIDE system (Nikon, Tokyo, Japan). Spatial transcriptomics was completed using the Visium Spatial Gene Expression system (10x Genomics, Pleasanton, CA, USA). Individual libraries were processed using 10x Genomics’ Space Ranger count pipeline. Length of the R2 sequences were limited to the first 50 bps. Space Ranger v2.1.0 count pipeline was used to manually align the fiducial frames of the Visium slide H&E images to gene expression data. For easier export, the Visium Space Ranger software down-samples the original high-resolution H&E image such that its longest dimension is 2000, resulting in the other dimension being resized based on the image’s original aspect ratio. The exported data frame of gene expression counts contains the Visium barcode coordinates from the original, full-resolution, H&E image. Due to the uniform down-sampling of the original image, a single scaling factor is provided by the software to match the original barcode coordinates to the dimensions of the exported, down-sampled, H&E image, to which the MPM images are registered.

### 2.4 Data Processing

All image registration was performed using the open-source TrakEM2 ImageJ plugin.^27, 28^ First, multi-channel MPM images were registered to one another in order to eliminate image drift between acquisitions, which could arise from variations in spectral properties of the system’s optics, physical movement of the tissue sample, and/or drift of the scanning mechanisms (Fig 2A). Following inter-channel registration, MPM images were then manually co-registered to the H&E reference image from the Visium dataset. The down-sampled H&E reference image matched to the Visium gene expression data was used as the final registration targets due to the original slide-scanned images being prohibitively large to process. Because the down-sampled H&E images are limited to 2000 pixels in the largest dimension, they were first scaled to dimensions roughly matching that of the MPM images. In the case of registering the MPM images to the Visium data, the down-sampled H&E image was scaled by a factor of three, changing its largest dimension to 6,000 pixels. With each MPM image having dimensions of roughly 5,500 x 5,500 pixels, any differences in scale were further accounted for during the registration process.

**Fig 2.**
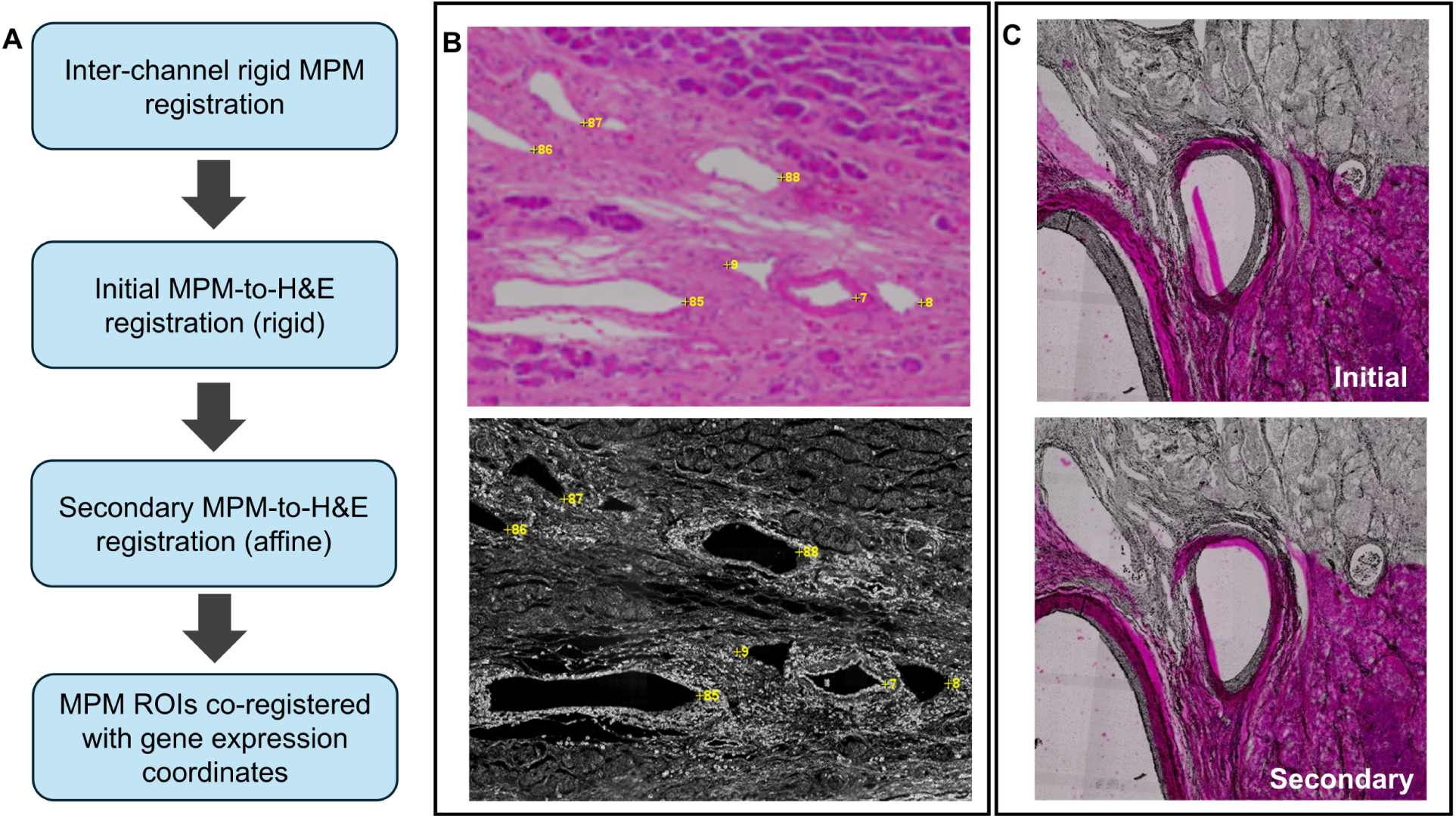
(A) Diagram of registration process to align the label-free image ROIs and transcriptomic signatures. (B) Large-scale morphological features were manually selected to register between H&E and MPM images. (C) Visual results of two-stage registration process.

For all subsequent registration steps, registration landmarks were manually placed on image regions that contained clear tissue landmarks, such as glands or cell clusters with unique morphology, as shown in Fig 2B. Landmarks were selected roughly uniformly over the full spatial area of each image and at least 80 landmarks were selected for each round. Multiple rounds of manual landmark placement and registration were necessary to register MPM images to the reference H&E image due to differences in scale and distortion of the tissue during the process of mounting it onto the Visium slide and H&E staining. An initial round of rigid registration through translation, rotation, and isotropic scaling was done to match image dimensions, and roughly over-lay landmarked regions. An affine transformation was then used with newly placed landmarks, repeating as necessary to align image contents (Fig 2C) to minimize mean-squared error. Following registration, tiles were visually compared between MPM and H&E to confirm accuracy. Registered images were then exported in their original 16-bit image format. The final result is the multi-channel MPM image being registered to a scaled version of the H&E image that defines the gene expression vector coordinates. The coordinates can be scaled by a factor of 3 (due to down-sampling) to locate the center of the field of view of the region of interest corresponding to a given gene expression vector (Fig 3).

**Fig 3.**
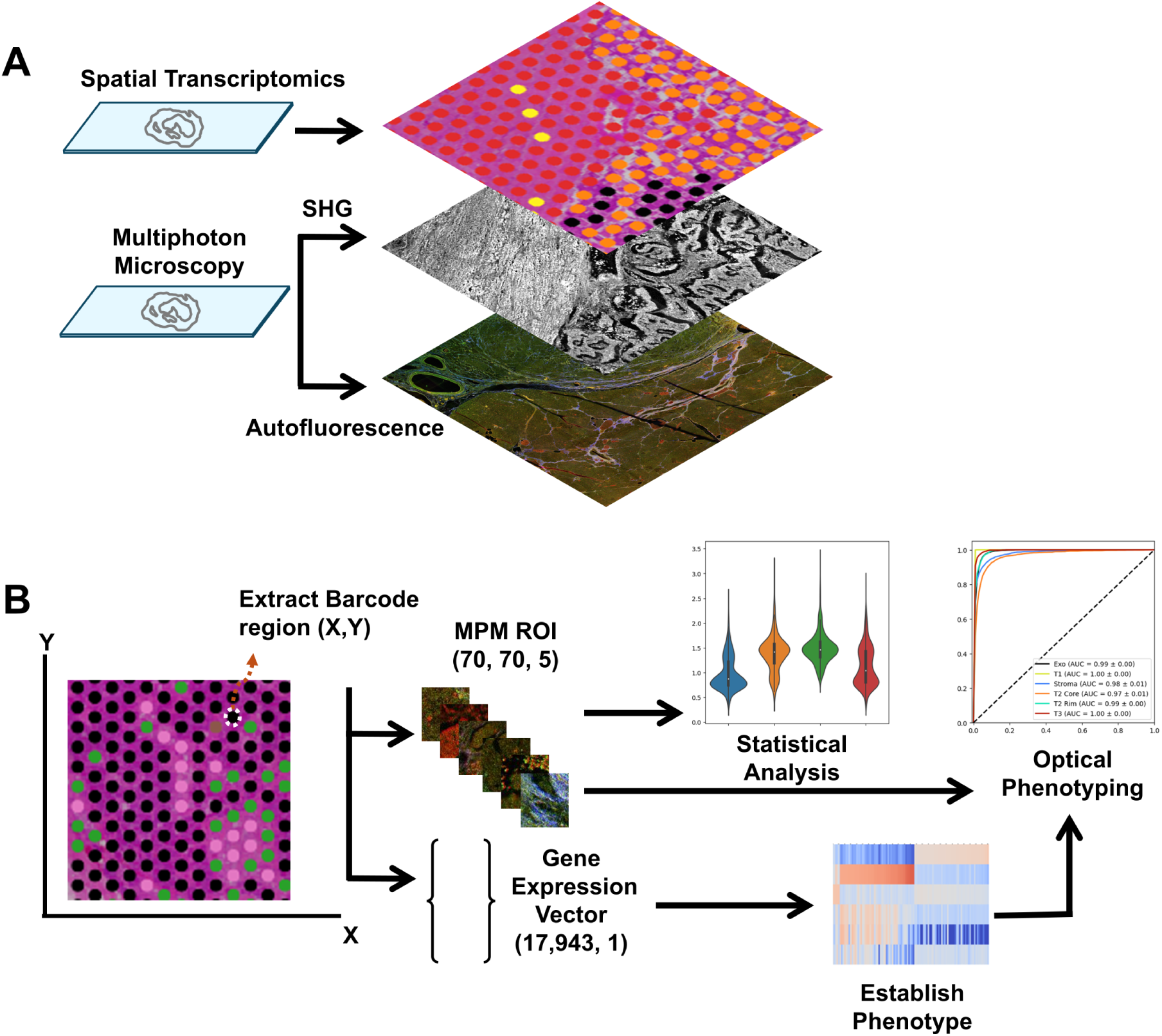
Schematic diagram of the study. (A) Using two adjacent patient sections, a co-registered dataset composed of spatial transcriptomics, autofluorescence, and SHG images was generated. (B) This dataset was then used to perform statistical analysis of imaging features between clusters defined by transcriptomic signatures, and to develop a deep learning model for optical phenotyping.

### 2.5 Statistical Analysis and Clustering

Gene expression data was imported into a Python environment as a data frame containing the unique molecular identifier counts for each human mRNA target and the pixel coordinates for the center of each barcode region. The pixel coordinates (defined in the space of the to the down-sampled H&E image) were then scaled by a factor of 3 to match the space of the registered MPM image. Regions of interest (ROIs) of each MPM image corresponding to the barcode pixel coordinates were extracted. ROIs were scaled by the pixel resolution of the MPM system to match the 100 microns spacing of the Visium barcode regions, yielding ROIs of approximately 70 x 70 pixels. A threshold was applied to the MPM ROIs to remove background (*<* 2000) and saturated pixels (*>* 65535) from any subsequent analysis. Abundance was measured by taking the mean value of each ROI corresponding to each channel. Channels 1,2,3 and 5 were normalized by dividing by the fourth channel in order to account for sample-to-sample fluctuations in light source intensity, working distance, and other systematic effects.

Tissue Phenotypes were established using k-means clustering of the transcriptomic signatures. Clustering of the spatial transcriptomic data was completed using the Loupe browser. K-means clustering was applied with k=6 and all four patient samples pooled together. The selection of k=6 was chosen by assessing the Silhouette score, as well as evaluating the spatial heterogeneity of cluster assignment. The results were then visualized by projecting each cluster onto the respective barcode region and superimposing this on an H&E brightfield image to visualize the clusters and spatial relationship (Fig. 4). Cell clusters were assigned tissue types and phenotypes by mapping gene expression profiles to the Human Protein Atlas and assessing the top 50 differentially expressed genes for each cluster as discussed in Section 3.1. Comparisons of MPM channels were performed for each cluster within a specific specimen, as well as for all tumor clusters separately. Data distributions are visualized using violin plots. The Shapiro-Wilk test was used to evaluate normality. All distributions were found to be significantly non-normal, and therefore, statistical significance was evaluated using the Kruskal-Wallis test followed by post-hoc Mann-Whitney U-

**Fig 4.**
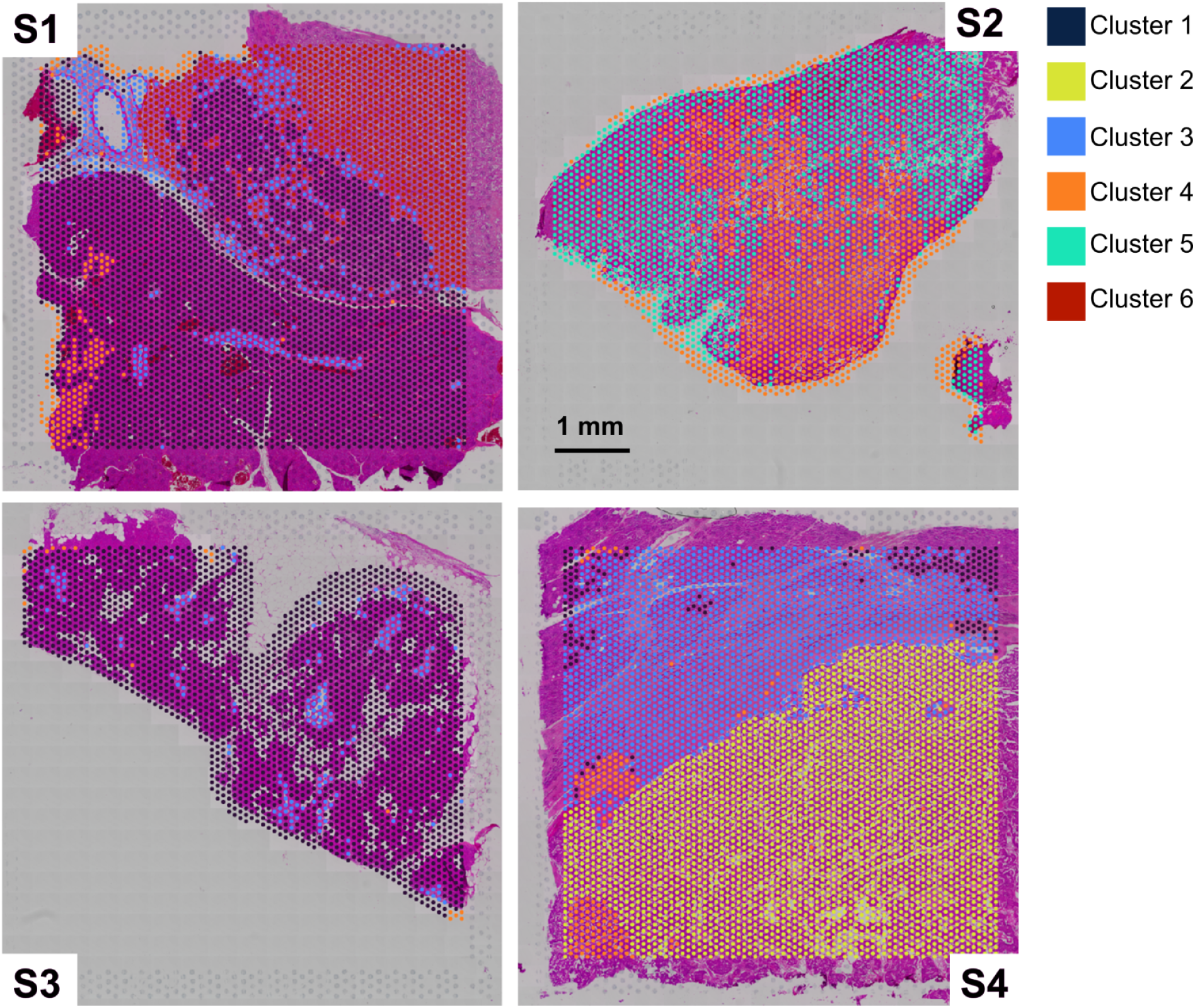
K-means clustering with k=6 performed for spatial transcriptomic data with all four patients pooled together. The clusters are superimposed on H&E images with each cluster positioned in the respective spatial barcode region and color coded depending on the cluster.

Test with multiple testing comparison using the Bonferroni correction method by multiplying the p-value by 60. For these comparisons, each ROI / barcode region is treated as a separate independent sample, which is in alignment with statistical analysis approaches in multi-omic studies using spatial transcriptomics.^29^

### 2.6 Image Classification

Three different classification approaches were developed to test the ability to perform optical phenotyping (Figure 5). We investigated these three different paradigms to understand what features within the image are most informative to predicting the tissue phenotype.

**Fig 5.**
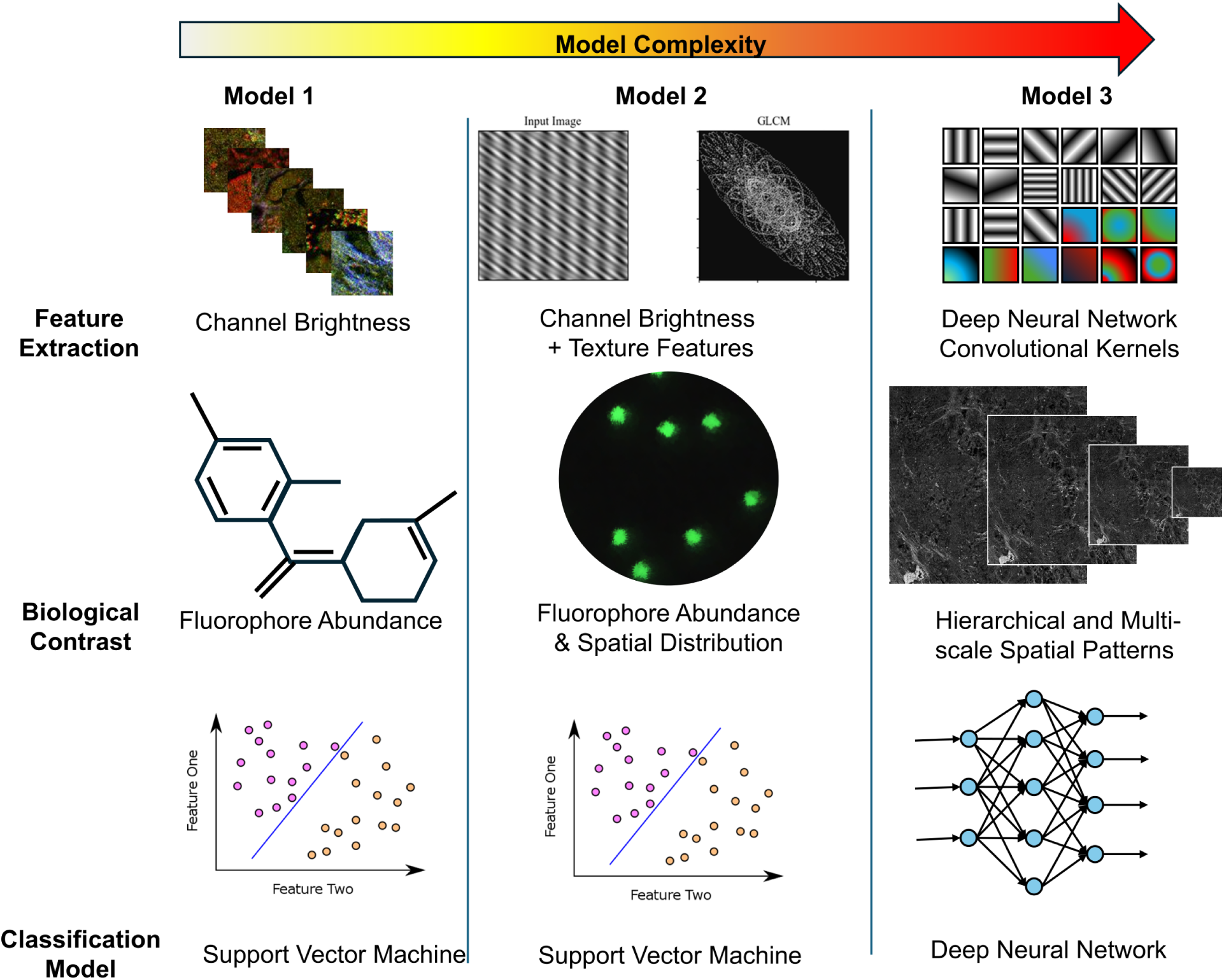
Overview of classification models evaluated in this study for optical phenotyping. Three models were assessed with increasing complexity. First, we tested support vector machine classifiers with average brightness of each channel used as input, followed by average brightness and image texture. Then we trained a deep learning model based on the VGG16 architecture.

For the first two models, we utilized classical machine learning with handcrafted features and a support vector machine. Our first model used only the average brightness of each channel, which is representative of the abundance of different contrast sources, yielding a total of 5 features. Our second model was trained with the original abundance features as well as features extracted using Haralick texture feature extraction,^30^ which quantifies the spatial relationship between adjacent pixel values, yielding a total of 70 features. Evaluating the spatial distribution of brightness is a measure of how the local microenvironment may be altered in the organization of different contrast sources, which is representative of the tumor microenvironment. For our third model, we utilized a convolutional neural network to perform unsupervised feature extraction. Here, the feature complexity increased further at the cost of explainability. However, this model can extract information at multiple spatial scales and incorporates non-linearities, which are likely necessary for modeling the complex sequence-function relationship in tissue.

For the first two models, the feature dataset was first standardized by centering at zero and scaling to unit variance for each channel. The support vector classifiers were trained with 80% of the dataset and the remaining 20% was used for testing. Five-fold cross validation with optimized weights was used for assessment for both scenarios, and all assessment metrics were averaged over all five folds.

For our third model, a customized convolutional neural network was designed to achieve high-accuracy multi-class image classification. Here, we applied transfer learning using a pretrained VGG16,^31^ an effective approach when limited labeled data is available, as it leverages features learned by the pretrained model. Figure 6A shows the customized VGG16 model architecture. The first four layers of the model were frozen to retain general feature representations extracted from the ImageNet dataset, while layers beyond the first maxpooling layer remained trainable to learn features specific to our MPM dataset. The batch size was set to 16. Additionally, a dropout layer was incorporated to mitigate potential overfitting, and early stopping monitor on the validation loss with patience of 6 epochs along with a learning rate scheduler was applied as part of the model’s callbacks to optimize training.

**Fig 6.**
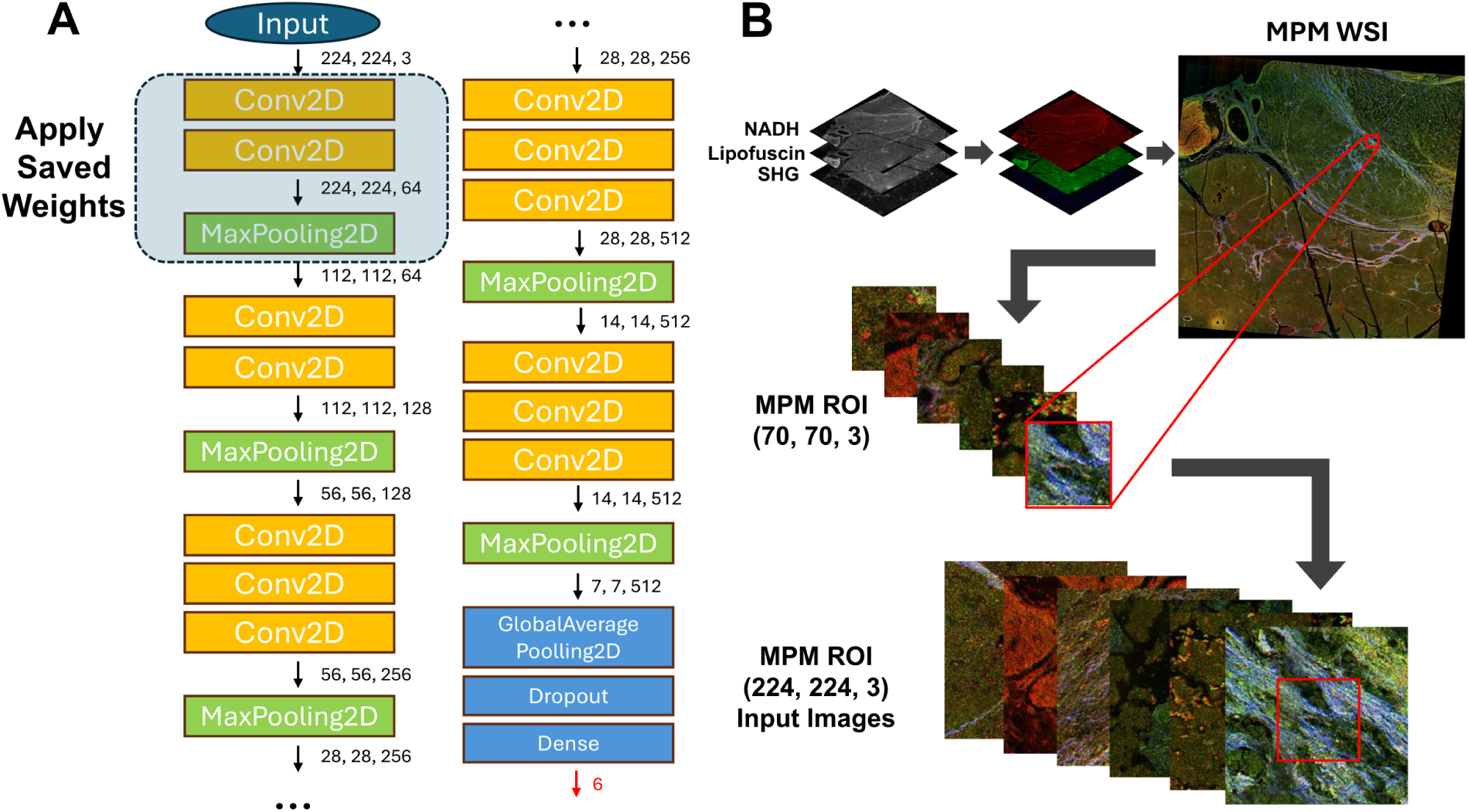
(A) Diagram of the customized pre-trained VGG16 architecture. The numbers next to the arrows represent the output shape of the preceding layer. The layers enclosed by the dashed line were frozen to preserve the pre-trained weights from the VGG16 model, originally trained on the ImageNet dataset. (B) Data preprocessing flowchart shows the RGB layering, tile extraction, and rescaling procedures for the input tile images.

Before the data was fed into the model, the four co-registered whole-slide MPM images were first converted into 3-channel RGB images to ensure compatibility with the network model as shown in Figure 6B. In this conversion, the red channel represented the NADH-dominant signal, the green channel corresponded to the lipofuscin-dominant signal, and the blue channel was assigned to the SHG signal. Based on gene expression data from each region of interest, tables were generated containing the coordinates of each region’s center along with their corresponding cluster labels. These colored MPM whole slide images (WSI) were then segmented into individual tiles using these tables. To effectively cover the entire barcode region, the ideal tile size for the MPM image was set to 70×70×3 as shown in Figure 6B. However, to match the model’s required input dimensions, each tile was resized to 224×224×3, incorporating additional surrounding areas of each barcode region. This re-sizing resulted in approximately 50% overlapping with adjacent regions of interest. Since neighboring areas generally share similar gene expression data due to high spatial correlation, this overlap did not introduce significant ambiguity to the model.

All images were then organized into separate folders, with 80% allocated for training, 10% for validation, and the remaining 10% reserved for testing to assess the model’s generalization. Additionally, subfolders were created to categorize images into different clusters. The “image dataset from directory” function from the utils library was used to efficiently load and preprocess the input images from the custom folder structure. All methods were implemented using python including sckit-learn, the TensorFlow deep learning framework and the Keras library on a PC equipped with an NVIDIA GeForce A4500 GPU. For all three classifiers, Receiver Operator Characteristic curves (ROCs) were generated for each class by evaluating the ability to classify each group in a “one vs. rest” paradigm. Confusion matrices were also generated to evaluate misclassification for each group in a true multiclass classification paradigm. Classification reports were generated that detail the precision, recall, and F1 score, as is standard for evaluating classification models. Together, these metrics provided a comprehensive assessment of the model’s performance. For this model, we utilized 10-fold cross validation to ensure all metrics reflected testing on the entire dataset. Visualization of the model’s classification performance was also visually evaluated by back-projecting incorrect classifications onto the location of the ROI on the original MPM images.

## 3 Results

### 3.1 Clustering and Definition of Phenotypes

Table 2 shows the results of K-means clustering with k=6 using the full transcriptomic data. Figure 7 shows the top 50 up- and down-regulated genes for each cluster. Of the six clusters, clusters 4 and 5 show the most similarity in gene expression, which is not surprising given that both are represented adjacent to one another in sample 2. Table 3 below lists key up-regulated genes for each cluster that is used for establishing tissue type and tumor phenotypes. Pathology reports were also evaluated for expression of specific hormones including glucagon and gastrin, as well as Ki-67 proliferative index.

**Fig 7.**
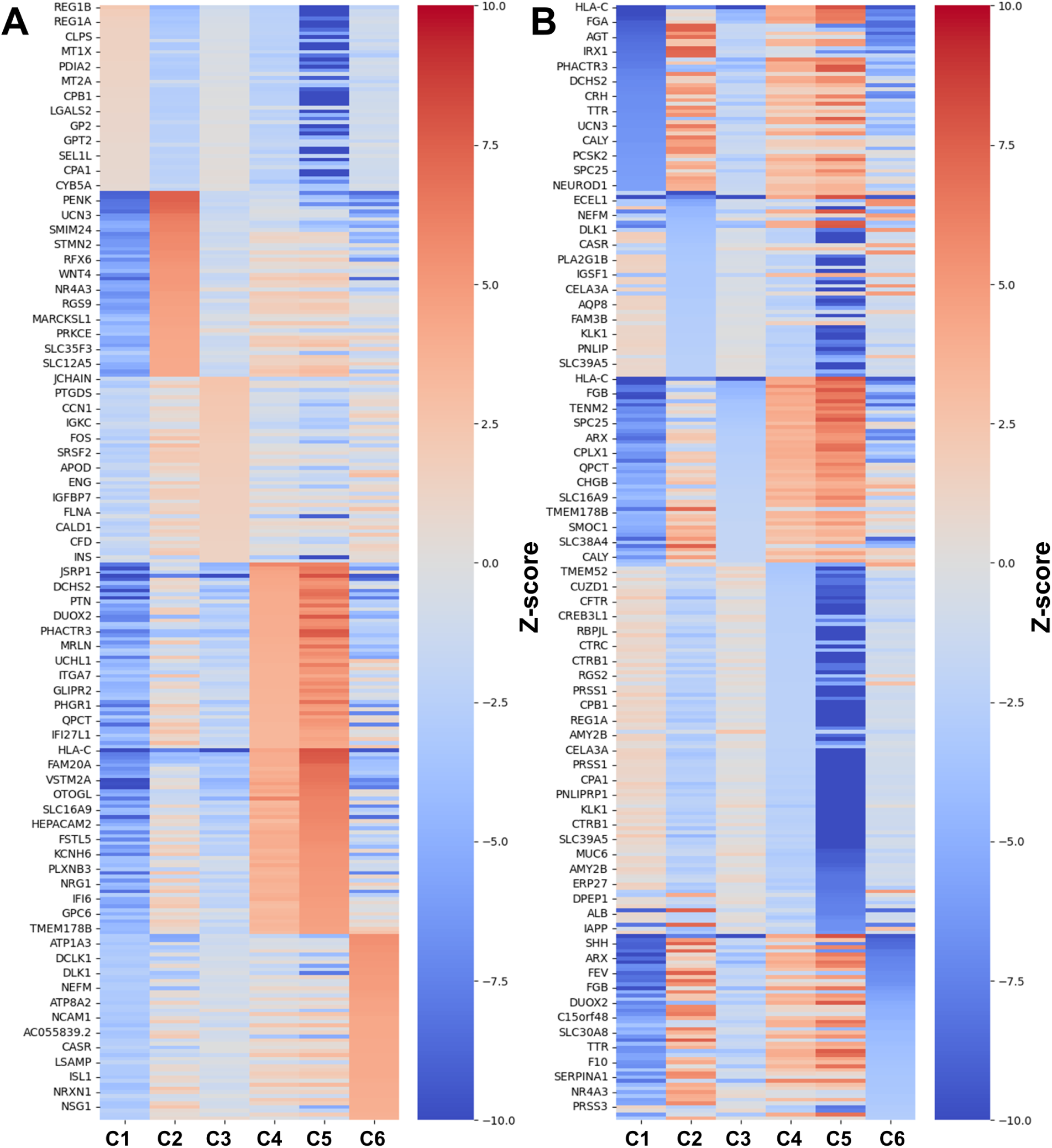
Gene expression heatmaps of the (A) top 50 over-expressed and (B) top 50 under-expressed genes for each cluster. Heatmap colors are denoted as the z-score comparing the expression levels of each gene for each cluster among the entire dataset.

**Table 2.**
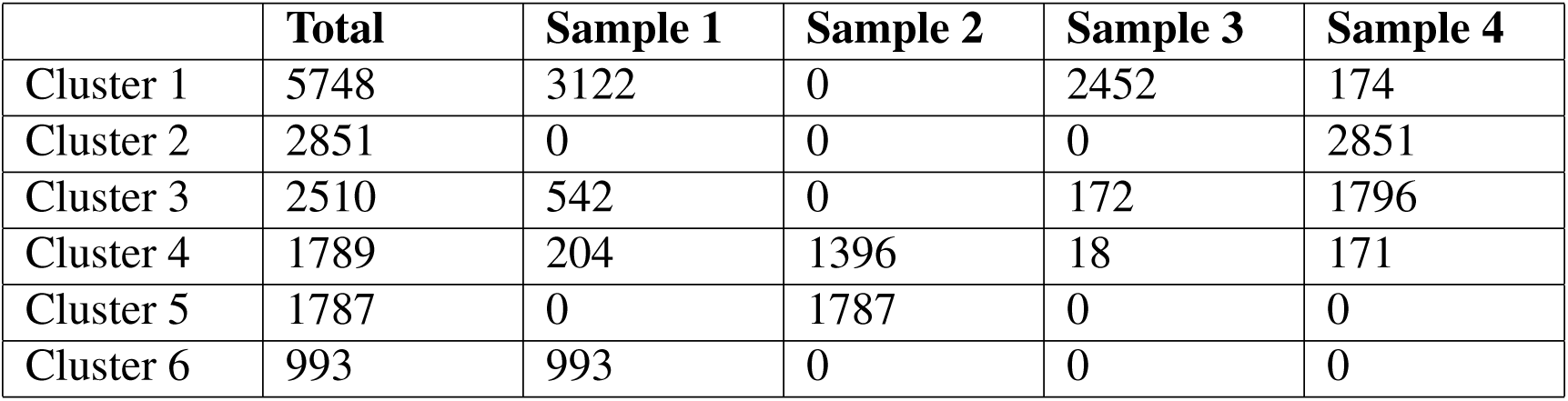
Results of K-mean clustering with k=6 using transcriptomic signatures. Numbers represent the total number of barcode regions assigned to each cluster across all samples.

|  | **Total** | **Sample 1** | **Sample 2** | **Sample 3** | **Sample 4** |
| --- | --- | --- | --- | --- | --- |
| Cluster 1 | 5748 | 3122 | 0 | 2452 | 174 |
| Cluster 2 | 2851 | 0 | 0 | 0 | 2851 |
| Cluster 3 | 2510 | 542 | 0 | 172 | 1796 |
| Cluster 4 | 1789 | 204 | 1396 | 18 | 171 |
| Cluster 5 | 1787 | 0 | 1787 | 0 | 0 |
| Cluster 6 | 993 | 993 | 0 | 0 | 0 |

**Table 3.**
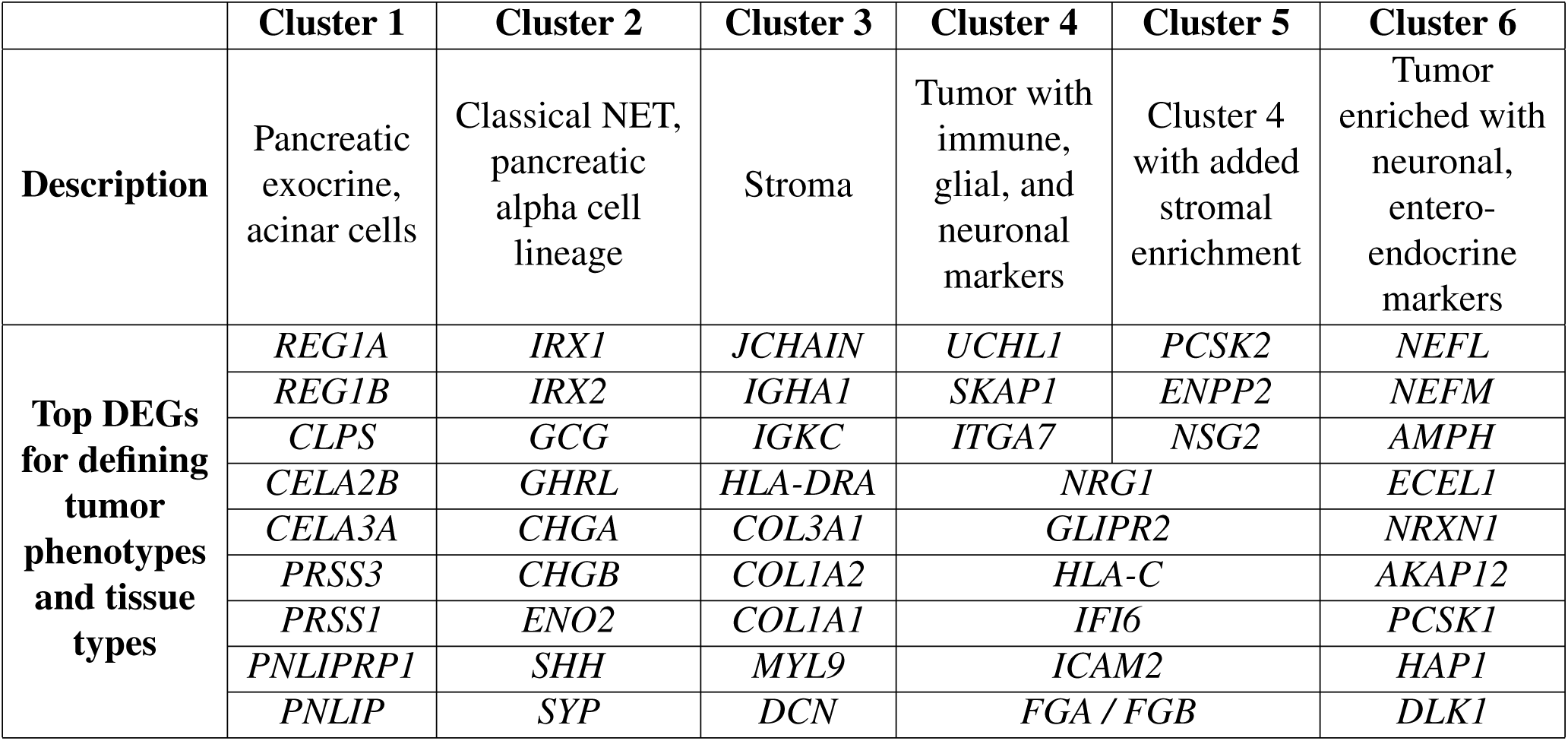
Interpreted description of each cluster and top differentially expressed genes (DEGs) for each used to establish phenotypes.

Cluster 1 is characterized by an enrichment in genes associated with acinar cells and the exocrine pancreas, including *REG1A/B*, *CLPS* and *CELA2B/3A* and *PNLIP*. This gene signature aligns with the expectation from pathological examination of the tissue.

Cluster 2 shows an enrichment in genes that are most classically associated with neuroendocrine tumors (e.g., *CHGA*, *CHGB*, *SYP*). Significant upregulated genes in this cluster include *IRX1*, *IRX2* and *GCG*, which are markers of the pancreatic alpha cell lineage. This is also consistent with the pathology report, showing positive tumor expression of glucagon. Interestingly, alterations of *CHGA* and *CHGB* at the protein level have been associated with different phenotypes in pancreatic neuroendocrine tumors; in this case, relating to aggressiveness.^32^ Tumor cells in this cluster also express the neuroendocrine tumor markers *SYP*, and *SHH*, a key developmental morphogen with a potential role in NET development.^33^ Therefore, cluster 2 is defined as a classical neuroendocrine tumor.

Cluster 3 exhibits upregulated expression of stromal and immune related genes. This includes an enrichment of plasma cell markers such as *JCHAIN*, *IGHA1*, and *IGKC*, immune cell markers including *HLA-DRA* and *COL3A1*, as well as markers of extracellular matrix and fibroblasts including *COL1A2*, *COL1A1*, *MYL9* (myofibrobalsts and smooth muscle) and *DCN* (endothelial cells). These markers align with the H&E image and pathological examination of the tissues. Thus, this cluster is defined broadly as stromal tissue.

Clusters 4 and 5 are occur within a single patient tumor and have similar transcriptomic profiles. However, clustering the dataset with n=5 clusters did not merge the two clusters, suggesting important differences between the two from the transcriptomic standpoint. Both clusters exhibit downregulated *PRSS1* expression, which is associated with hereditary pancreatitis,^34^ as well as *NRG1*, *GLIPR2*, *HLA-C*, which are immune response-related markers, suggesting that the tumor may exhibit unique immune or inflammatory characteristics. Other shared features include expression of *NRG1* and *GLIPR2*, markers of glial cells, as well as *FGA* and *FGB*, which are related to the blood clotting response. Important differences between clusters 4 and 5 are the expression of *ENPP2* in cluster 5 (a stromal marker) in addition to a seemingly lower expression of genes for cluster 4 universally. This may suggest that cluster 4 represents some loss of gene expression, perhaps due to necrosis. As such, we define cluster 4 and 5 as a tumor with immune features, with 5 being denoted “rim” and 4 being noted “core,” where some necrosis may be present. Clinically, this was the only specimen that was reported focally positive for gastrin expression, which is often related to a specific subtype of pancreatic NET.^35^ Furthermore, the tumor specimen relating to clusters 4 and 5 yielded a clinical Ki-67 index of 8%, which was double the corresponding index of the specimens associated with clusters 2 and 6 (both of which were measured around 4%).

Finally, cluster 6 exhibits enrichment of enteroendocrine markers (*PCSK1*, *HAP1*, *DLS1*), as well as neuronal markers, including the expression of *NEFL*, *NEFM*, *NCAM1*. Distinct differences are observed between cluster 6 and 2, and notably this cluster is associated with a tumor that was reported as glucagon negative. Additionally, there are a number of prognostic markers such as over-expression of *SYT17* and under-expression of *SHH*, which in other pancreatic cancers at the protein level have been associated with more favorable prognosis.^36^ As such, we define cluster 6 as tumor with enteroendocrine and neural features. Referencing clinical data, the patient in which this specimen was observed was the only patient who possessed lymph node metastases – which is highly correlated with poor survival.^37^

#### 3.1.1 Statistical Analysis of Image Features

Figure 8 shows the correlation between imaging channels before and after normalization. Prior to normalization by the porphyrin channel, interchannel correlation was high, over 0.65 on average. The SHG is lower in general, which is expected given that the optical process is not a fluorescence process. After correlation, the correlation is significantly reduced, being below 0.3 on average for all pairs of channels.

**Fig 8.**
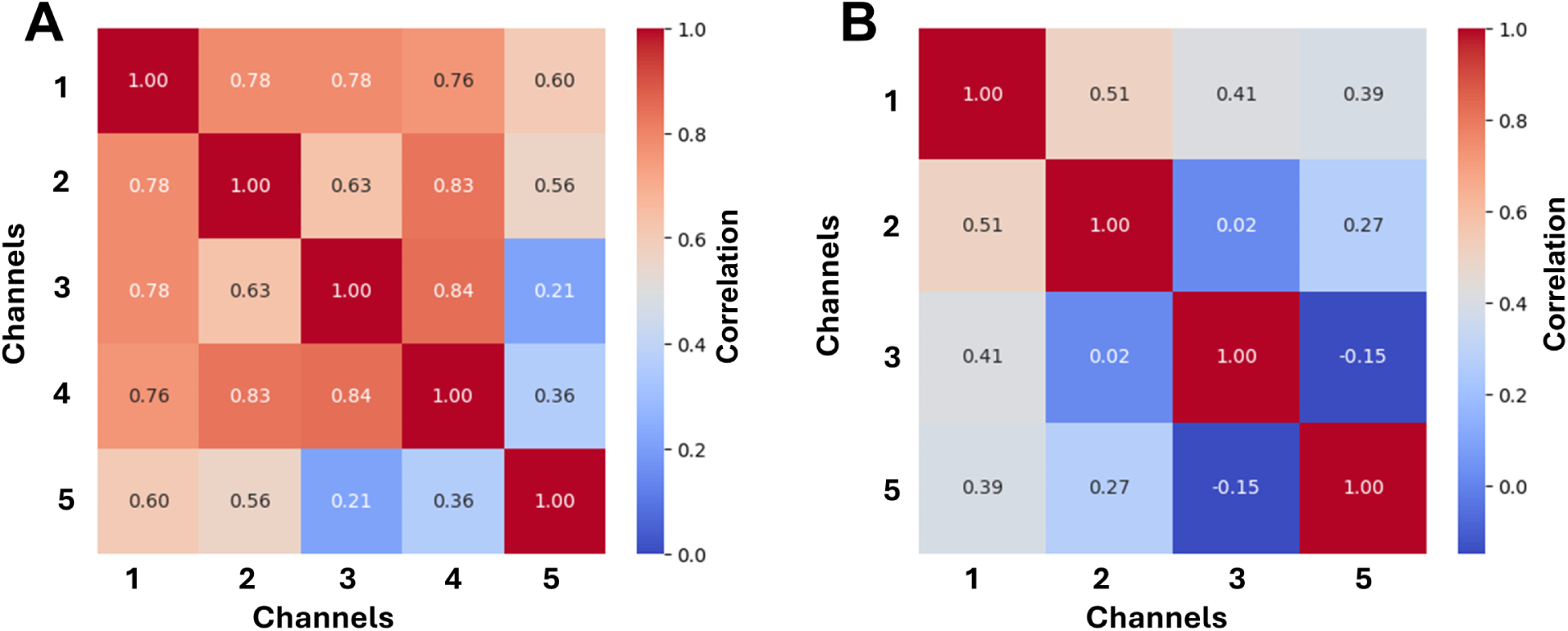
Inter-channel correlation of MPM imaging channel s (A) before and (B) after normalization by channel 4.

Figure 9 shows the violin plot comparing normalized average normalized brightness of each channel for ROIs assigned to different clusters. Significance is denoted above each cluster as *** for p*<*0.001 for all pair-wise comparisons except the pairs denoted with a line showing “n.s.” for not significant. Several observations can be made. First, each cluster exhibits unique distribution across the four channels. For example, while channel 1 distribution is similar for the first three clusters (exocrine, T1 and stroma), the distributions differ significantly in channels 2 and 3. This suggests that the multi-channel information is important for successfully differentiating tissue types. Among all clusters, the two clusters from T2 – the rim and the core – show the most similar distributions, which is supported by the lack of statistical significance in two of the channels. There are subtle differences between the two, for example a proportion of the T2 Core cluster is contained in a lower tail of the distribution (indicating lower brightness, potentially related to necrosis). Given that both clusters are closely related transcriptomically, the similarity of the imaging characteristics is not surprising. This also indicates that the two clusters may be difficult to differentiate during classification. Considering this data overall, while most clusters have clear differences in the distributions among one another (supported by the strong statistical significance), there are still significant overlaps in the image brightness distributions. The consequence of this is that there likely will be significant challenges classifying clusters based on this information alone, motivating the need for more advanced image features such as texture and other spatially-derived features.

**Fig 9.**
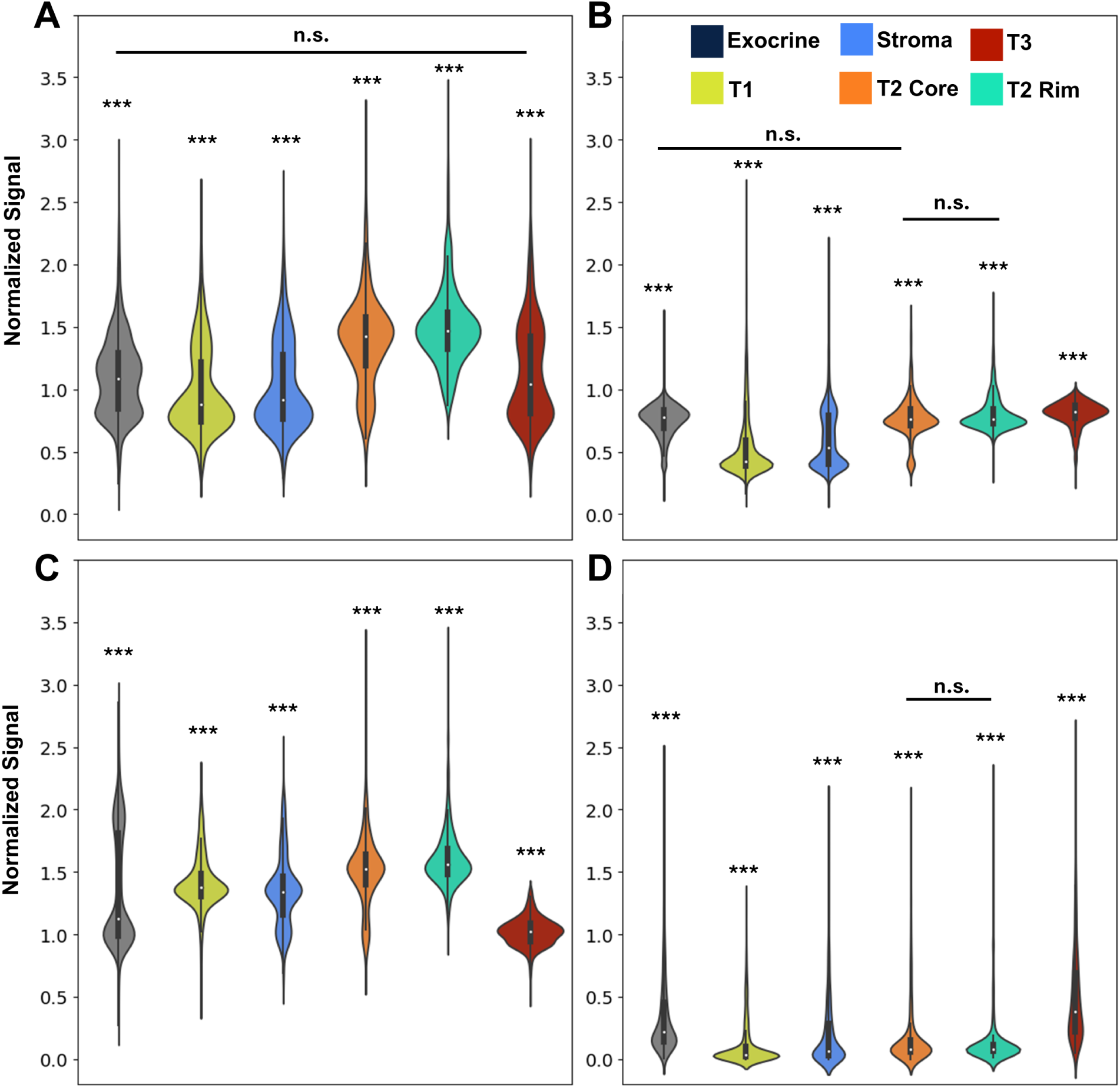
Violin plots comparing average brightness for ROIs assigned to different tumor clusters for normalized (A) channel 1, (B) channel 2, (C) channel 3, and (D) channel 5. Significance is denoted *** for p<0.001 for all pair-wise comparisons. Bars denoting no significant difference are used to show pairs that are not significantly different.

### 3.2 Image Classification

The classification results for each of the three models are shown below in Tables 4-6, which present the recall, precision, and F1 score for each class and classifier, respectively, averaged over all five folds. Included in these tables is the AUC for the one-vs-rest classifiers shown later. For the two support vector classifiers, the overall performance is marginal. The classifier trained on abundance only (Table 4) is unable to classify class 3 and 4 – this most likely is due to a lack of sufficient features in the model, as only 5 channels are present, whereas 6 classes exist. The overall weighted average accuracy of the classifier was 43.8% ± 0.83%.

**Table 4.**
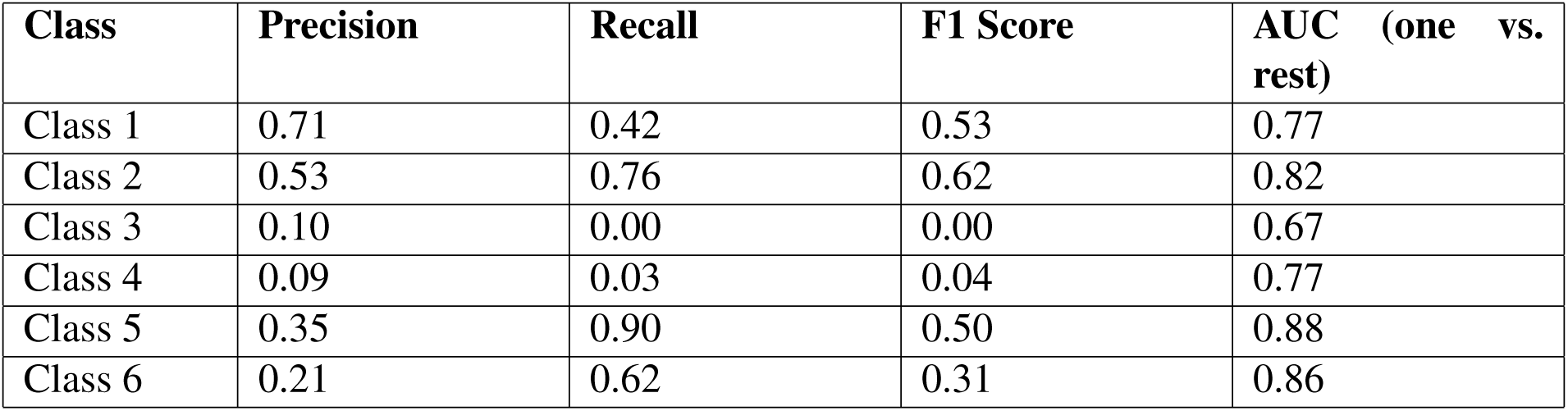
Precision, recall, and F1 score for support vector classifier trained on abundance, averaged across all five folds.

| Class | Precision | Recall | F1 Score | AUC (one vs. rest) |
| --- | --- | --- | --- | --- |
| Class 1 | 0.71 | 0.42 | 0.53 | 0.77 |
| Class 2 | 0.53 | 0.76 | 0.62 | 0.82 |
| Class 3 | 0.10 | 0.00 | 0.00 | 0.67 |
| Class 4 | 0.09 | 0.03 | 0.04 | 0.77 |
| Class 5 | 0.35 | 0.90 | 0.50 | 0.88 |
| Class 6 | 0.21 | 0.62 | 0.31 | 0.86 |

**Table 5.** Precision, recall, and F1 score for support vector classifier trained on abundance and image texture features, averaged across all five folds.

| Class | Precision | Recall | F1 Score | AUC (one vs. rest) |
| --- | --- | --- | --- | --- |
| Class 1 | 0.78 | 0.74 | 0.76 | 0.91 |
| Class 2 | 0.57 | 0.45 | 0.51 | 0.92 |
| Class 3 | 0.36 | 0.39 | 0.37 | 0.76 |
| Class 4 | 0.37 | 0.21 | 0.26 | 0.81 |
| Class 5 | 0.52 | 0.67 | 0.58 | 0.93 |
| Class 6 | 0.22 | 0.41 | 0.28 | 0.86 |

**Table 6.**
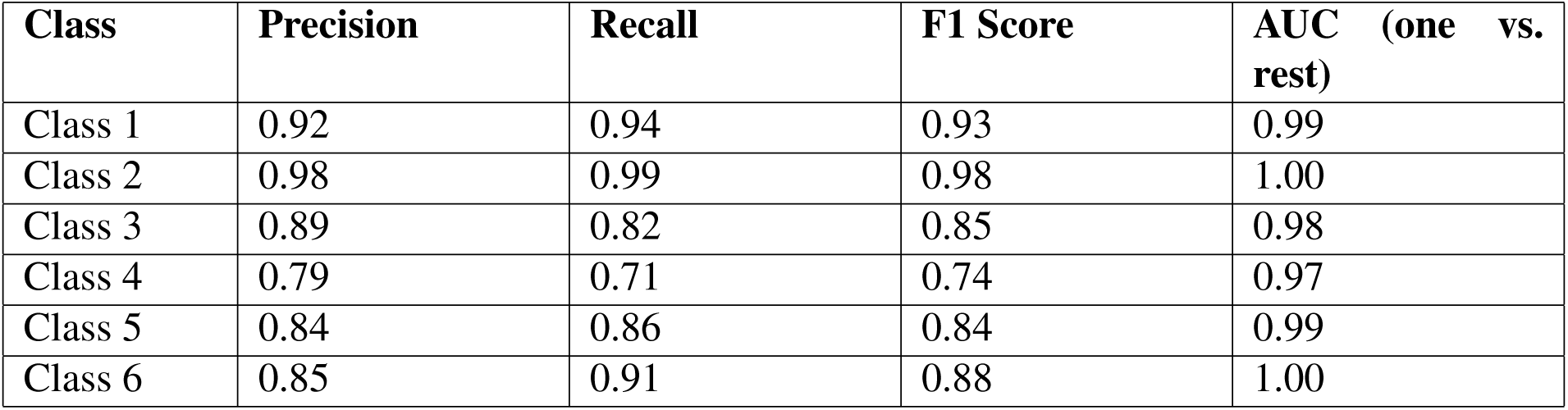
Precision, recall, and F1 score for the deep neural network model averaged across all ten folds.

For the support vector classifier trained on image texture and abundance (Table 5), there is moderate improvement. The average weighted classification accuracy increases to 54.4% ± 0.54% overall, and performance on a class-by-class basis increase nearly universally with the exception of a slight decrease in performance for class 2. Classes 4 and 6 remain the most challenging to classify accurately.

In stark contrast to the support vector classifiers with handcrafted features, using deep learning for feature extraction and classification, shown in Table 6, yields significant improvements, with an overall weighted classification accuracy of 89.4% ± 1.5%. The misclassification Class 4 is the primary factor affecting overall prediction accuracy, which also performed poorly for the two prior classifiers.

Visualizations of ROC curves and confusion matrices for all classes are shown for each classifier in Figure 10. Figure 10A-C presents the one-vs-rest ROC curves for all classes, highlighting the varying classification performance across different categories. As observed in the plot and tables, the AUC values are consistently higher than the F1 scores across all classes, as AUC evaluates the model’s ability to distinguish between classes across a range of decision thresholds. Even if a suboptimal decision threshold results in classification errors and lower F1 scores, the model may still exhibit strong discriminative capability over the entire probability distribution. The ROC curve from the deep neural network model exhibits a sharp ascent toward the top-left corner, particularly for the T1 and T3 classes. This behavior reflects a strong balance between sensitivity and specificity, effectively minimizing false positives while maintaining a high true positive rate. For the simpler models using handcrafted features, the AUC is relatively high for a subset of classes (e.g. T2 Rim and T3), but relatively low for other classes (Stroma, T2 Core).

**Fig 10.**
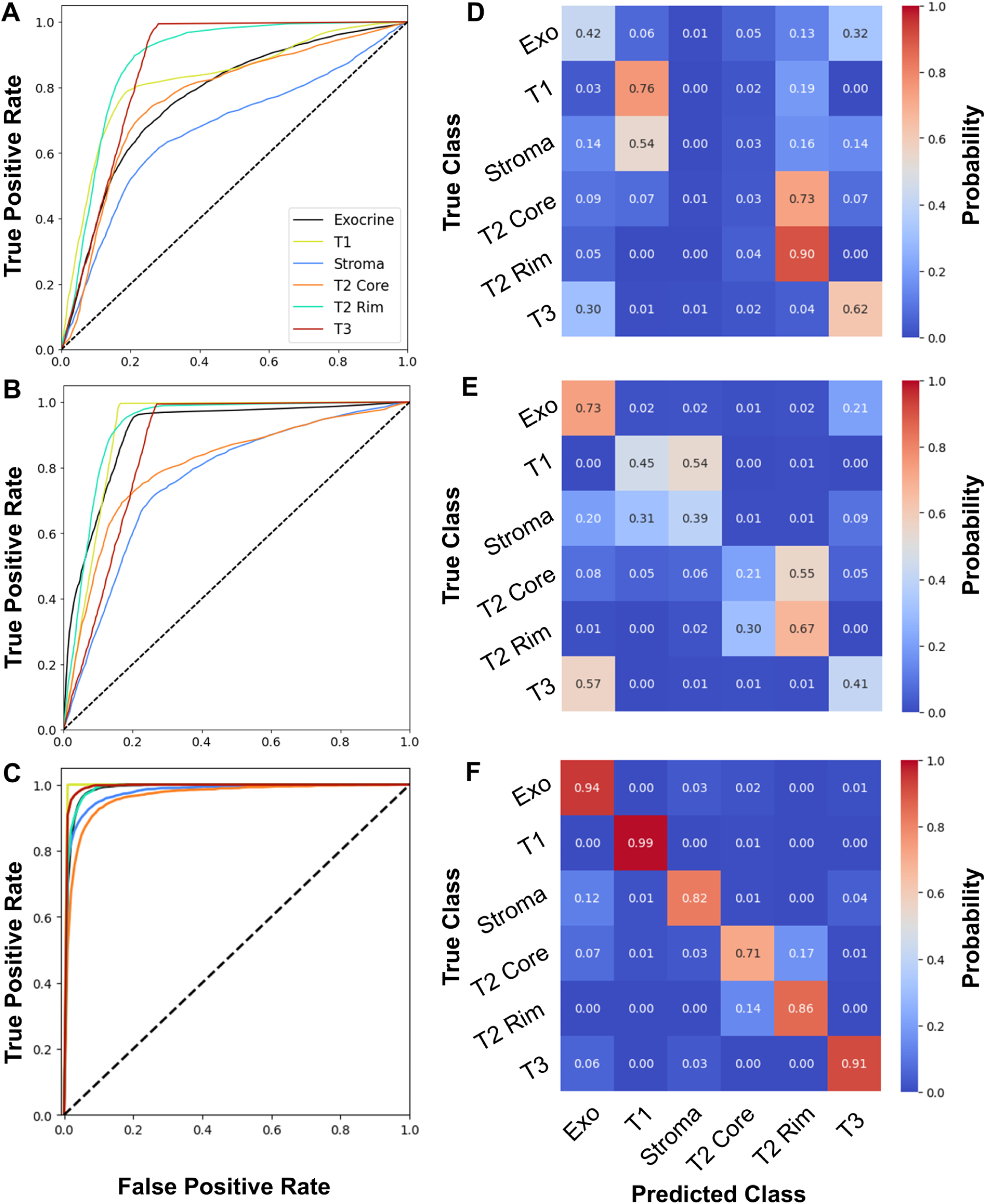
Classification model results. ROC curves to assess one-vs-rest classification performance for (A) support vector machine with abundance features, (B) support vector machine with abundance and texture, and (C) deep neural network. Confusion matrices for multi-class classification for (D) support vector machine with abundance features, (E) support vector machine with abundance and texture, and (F) deep neural network.

The issue of misclassification is most clearly illustrated in the confusion matrices (Figure 10D-F). For the two support vector classifiers, the models consistently misclassify T1 and stroma, as well as T2 rim and core. The latter is not surprising given the earlier observations of the similarities between those two clusters both transcriptomically and optically. For the deep learning model, the ability to differentiate all classes is drastically improved; however, misclassification between exocrine and stroma persists, along with a notable number of misclassifications between the T2 core and rim. The latter two classes were also commonly misclassified in the support vector approach. This aligns well with our discussion of the gene expression and phenotype clustering, as these two classes share similar transcriptomic profiles and are both highly represented in a single tissue specimen. The misclassification of stroma and exocrine is likely due to the fact that the ROIs contain a multitude of cell types and therefore are not “pure.” Some may contain both exocrine and stromal cells, resulting in some reduction in image classification performance.

Figure 11 presents a back-projection of all misclassified ROIs onto MPM WSI using the deep learning model from the test dataset across the 10-fold models, providing a detailed visualization of recurring misclassification patterns. By mapping these instances, we can identify spatial and structural trends contributing to classification errors, revealing potential model biases and overlapping feature distributions among classes. As illustrated in the legend patch on the third MPM WSI (Figure 11C), the inner circle color of each individual spot represents the predicted class, while the outer circle indicates the true class. Across the four WSIs, the majority of mixed-color groups correspond to blue-black (Exo and Stroma) and cyan-orange (T2 rim and core), aligning with our previous analysis.

**Fig 11.**
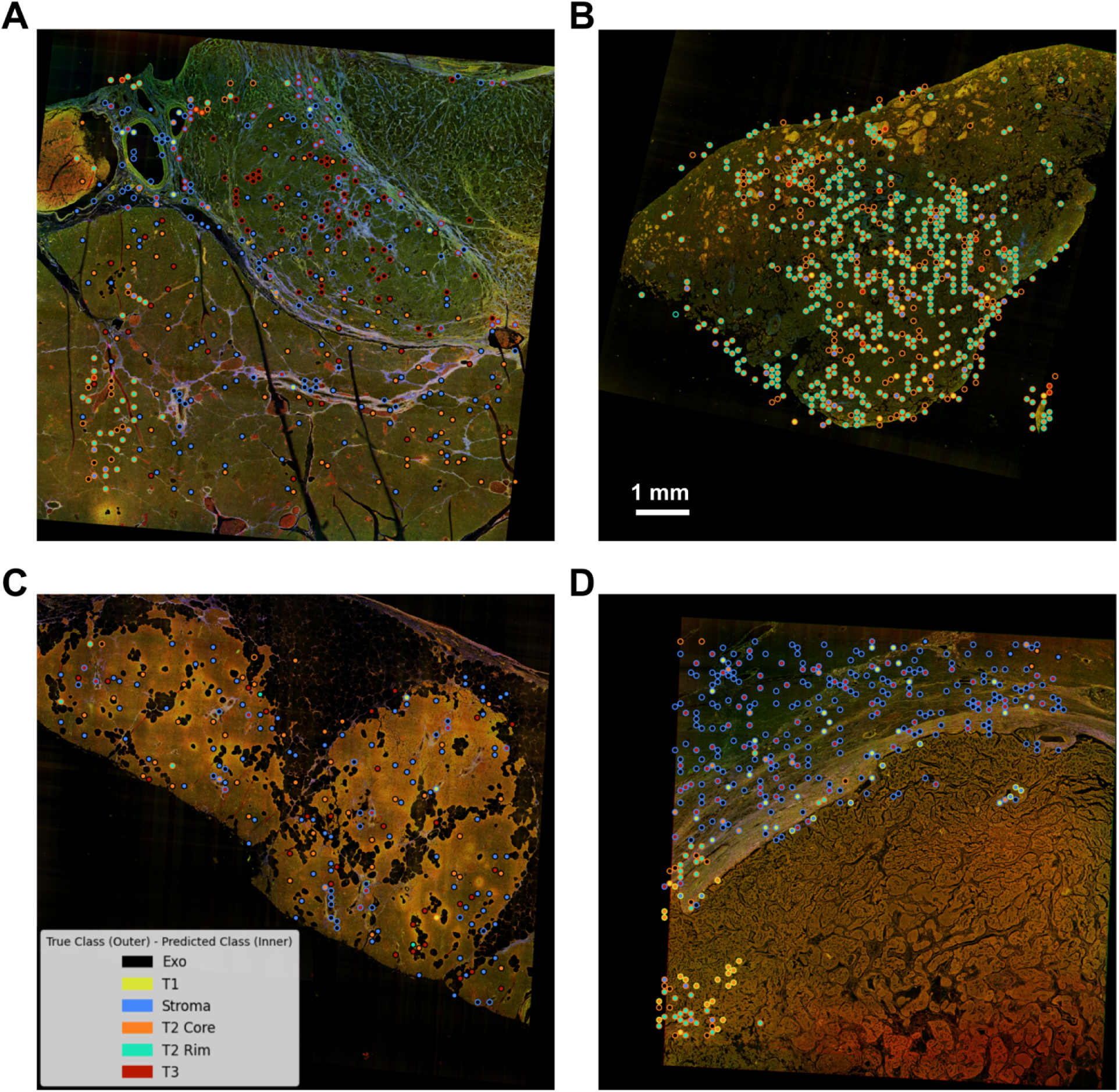
Back projection of all misclassified ROIs from the test dataset across the 10-fold models onto the four wholeslide images provides a comprehensive visualization of recurrent misclassification patterns. The color of the center of the circle represents the predicted class whereas the color of the rim represents the true class.

Notably, the second MPM WSI represents an all-tumor slide, containing both T2 core and T2 rim classes. In Figure 11B, most misclassified images cluster within the T2 core region, with errors predominantly occurring at the interface between two clusters. This misclassification may partially stem from the expanded input image size, where adjacent tiles share overlapping regions, potentially introducing bias, particularly at class boundaries. However, given the limited dataset, the benefits of this expansion outweigh the loss introduced by overlapping biases. Furthermore, T1 (Figure 11D, lower right region) and T3 (Figure 11A, upper right region) exhibit an exceptionally high true positive rate, with nearly no false predictions. There are also some misclassified instances that lie outside the WSI region, indicating imperfections in data preprocessing that may introduce additional bias.

These observations suggest potential improvements to enhance model performance. Refining the data preprocessing pipeline by implementing a more precise tile screening method would help accurately exclude dark images and irrelevant tiles, reducing unnecessary biases. Additionally, once more data becomes available, limiting the overlap between adjacent tiles could minimize misclassification at class boundaries. Another possible refinement involves developing specialized sub-models for Exo vs. Stroma and T2 core vs. rim, which may improve classification accuracy for these challenging class pairs. By addressing these factors, the model’s robustness and predictive performance can be further improved.

Figure 12 presents complementary plots to Figure 11, showing all the correct predictions backprojected onto the four WSIs. These two figures map all the prediction results back onto the MPM WSIs, showing consistency with the spatial transcriptomics profiles. Ultimately, the strong performance on the ROC curves, coupled with the high overall accuracy (89.4%) on the test dataset, further validates our deep learning model’s predictive capability and reinforces the potential of integrating label-free microscopy and spatial transcriptomics as a powerful tool for optical phenotyping.

**Fig 12.**
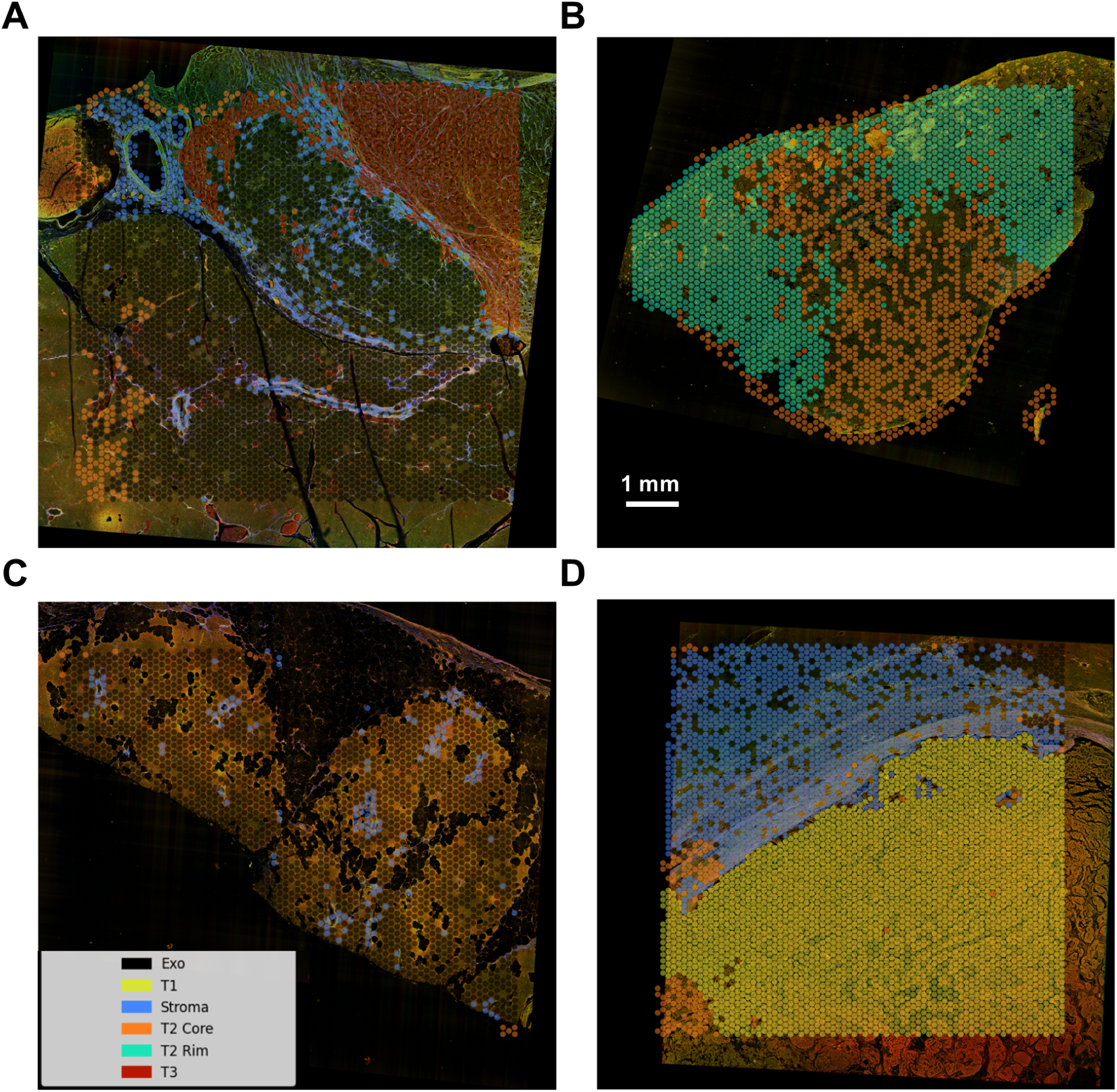
Back projection of all correct predicted ROIs from the test dataset across the 10-fold models onto the four whole-slide images

## 4 Discussion

The primary goal of this work was to develop a method for optical phenotyping of tissues using label-free microscopy. The results of this work demonstrate several important findings. First, label-free optical imaging contrast, in this case related to autofluorescence and second harmonic generation, carries significant biologically relevant information that can be used for optical phenotyping. The evaluation of different feature extraction methods and classification approaches supports our hypothesis that multi-scale hierarchical features are necessary to more comprehensively capture the tissue microenvironment and enable optical phenotyping.

Despite our promising findings, there are limitations that must be acknowledged and addressed in future work. First, this work is carried out using *ex vivo* FFPE specimens, and the fixation process is known to alter tissue autofluorescence. This limits our ability to draw specific conclusions about what fluorescence species are varying in abundance, as there is significant disagreement in the literature regarding what fluorescence species are preserved.^38, 39^ In addition, while the approach of using label-free imaging for in vivo optical phenotyping is theoretically possible, all findings will need to be validated and tested in fresh / *in vivo* tissues given the differences in image content compared to FFPE. Likely, the features learned by the DNN using fresh tissues will differ significantly from FFPE, indicating that a different model would need to be developed.

Similarly, our dataset was limited to only four specimens due to both cost and availability of specimens that yielded sufficient RNA quality. While the number of barcode regions numbered close to 15,000 total, the small sample size in terms of independent patients required us to segment the dataset treating each of the barcode regions as independent. While this is common in multiomic studies, a more rigorous approach with a larger dataset and more patient specimens is required to fully evaluate the statistical properties and generalizability of our classification models.

Another important discussion is the clustering approach to define our phenotypes. The approach presented in this study involved a completely unbiased and unsupervised method to reduce any subjective interpretation and selection of phenotypes. While powerful for maintaining reproducibility and rigor, the approach has some inherent limitations. The mathematical assignment of clusters using k-means clustering does not necessarily integrate specific biological information such as spatial correlation of the signatures, and weighting of different genes that may have greater significance. This may lead to some tissues being clustered incorrectly due to numerical noise. Indeed, the clustering of our specimens seemed to suggest there was a small degree of this; for example, cluster 4 appearing in the normal tissue region of specimen S1 (See Figure 4). It is also important to recognize that there are many more than 6 cell types present overall. The spatial sampling for transcriptomics used in this study was limited to spacing the sampling points by 100 microns; thus within each cluster, there could be a variety of cell populations contributing to the overall transcriptomic signature, which will result in a wider distribution within each cluster. Single cell sequencing would limit this effect, but would also lose the surrounding microenvironment information given by the larger field of view of our approach. One of the most significant outcomes of this work is that our results suggest that the local environment is critical to effectively performing optical phenotyping. Furthermore, our phenotyping is focused on tissue-level properties, not cell-level properties, therefore single cell sequencing is likely not appropriate for measuring these effects. Recently, the technological platforms for spatial transcriptomics have been advanced to enable higher resolution spatial sequencing over what was used in this study. Integrating these next generation technologies would be a valuable way to assessing phenotypes with higher precision without losing local environmental characteristics.

Finally, integrating a deep learning approach requires future work investigating the explainability of the model to determine specifically what features are used for performing high accuracy classification and to evaluate whether these features are generalizable to other tissues and patients. In addition, an interesting avenue of future inquiry is to investigate how the transcriptomic signatures could be analyzed to assess higher level biological processes such as pathway regulation. This information could be used to inform and validate what image features offer the most predictive value. Along these lines, other label-free imaging modalities such as polarization and fluorescence lifetime imaging could be integrated into this architecture and evaluated for their ability to perform optical phenotyping. Ultimately, the framework we present in this manuscript has wide potential for addressing a number of intriguing scientific questions and lays the foundation for accelerating discoveries in the field of biophotonics.

## 5 Conclusion

Phenotyping is a powerful tool in diagnostics and for developing precision therapeutics, but most phenotyping methods are costly, time intensive, and not compatible with *in vivo* use. In this study, we introduce a new approach for “optical phenotyping” using label-free microscopy images, deep learning, and tissue phenotypes as defined by spatial transcriptomics. Our results show significant promise for optical phenotyping. The presented method is able to achieve over 89% accuracy in a complex multi-class classification problem. While promising, a number of outstanding challenges are to be addressed in future work; namely, expanding the patient cohort, addressing non-biological heterogeneity that may influence phenotype assignment, as well as eventual testing in fresh tissues.

## 7 Code and Data Availability Statement

All code and data associated with this work will be made available upon publication.

## 8 Acknowledgements

We would like to thank Carole Kepler and Alexandra Jimenez and the Tissue Acquisition and Cellular/Molecular Analysis service or assistance with histological processes, as well as Martin Deymier at the University of Arizona Genetics Core. We would also like to thank the UC Davis Comprehensive Cancer Center for supporting this work.

**Shuyuan Guan** is currently a PhD candidate at the University of Arizona’s Wyant College of Optical Sciences. She works in the “Biomedical Optics and Optical Measurement” Laboratory with Dr. Travis Sawyer. Her research interests involve the design and development of optical imaging systems, and leveraging deep learning techniques to enhance the outcomes of biomedical imaging.

**Travis Sawyer** is an Assistant Professor of Optical Sciences University of Arizona’s Wyant College of Optical Sciences. His laboratory focuses on developing novel optical technology for healthcare applications including cancer screening, minimally-invasive procedures, and biomarker identification, with a particular emphasis on gastrointestinal cancer.

## References

1 R. Wang and Z. Wang, “Precision medicine: Disease subtyping and tailored treatment.,” Cancers (Basel). 15(15), 3837 (2023). [doi: 10.3390/cancers15153837.].

2 A. Sahu, K. Kose, L. Kraehenbuehl, et al., “In vivo tumor immune microenvironment phenotypes correlate with inflammation and vasculature to predict immunotherapy response.,” Nat Commun. 13, 5312 (2022). [doi:10.1038/s41467-022-32738-7].

3 C. Torres and P. Grippo, “Pancreatic cancer subtypes: a roadmap for precision medicine.,” Ann Med. 50(4), 277–287 (2018). [doi:10.1080/07853890.2018.1453168].

4 H. G. Weng C, Shah NH, “Deep phenotyping: Embracing complexity and temporality-towards scalability, portability, and interoperability.,” J Biomed Inform. 105, 103433 (2020). [doi:10.1016/j.jbi.2020.103433].

5 P. Harrison, A. Wright, and J. Mank, “The evolution of gene expression and the transcriptome-phenotype relationship.,” Semin Cell Dev Biol. 23(2), 222–229 (2012). [doi:10.1016/j.semcdb.2011.12.004].

6 N. Shaked, S. Boppart, L. Wang, et al., “Label-free biomedical optical imaging.,” Nat. Photon. 17, 1031–1041 (2023). 10.1038/s41566-023-01299-6.

7 A. Croce and G. Bottiroli, “Autofluorescence spectroscopy and imaging: a tool for biomedical research and diagnosis.,” Eur J Histochem. 58(4), 2461 (2014). [doi:10.4081/ejh.2014.2461].

8 T. Knapp, S. Duan, J. Merchant, et al., “Quantitative characterization of duodenal gastrinoma autofluorescence using multiphoton microscopy.,” Lasers Surg Med. 55(2), 208–225 (2023). [doi:10.1002/lsm.23619].

9 A. T. Shah, T. M. Heaster, and M. C. Skala, “Metabolic imaging of head and neck cancer organoids,” PLOS ONE 12, e0170415 (2017). 10.1371/journal.pone.0170415.

10 J. Bec, D. Vela, J. Phipps, et al., “Label-free visualization and quantification of biochemical markers of atherosclerotic plaque progression using intravascular fluorescence lifetime,” JACC Cardiovasc Imaging. 14(9), 1832–1842 (2021). [doi:10.1016/j.jcmg.2020.10.004].

11 N. Ghosh and I. Vitkin., “Tissue polarimetry: concepts, challenges, applications, and outlook,” J Biomed Opt. 16(11), 110801 (2011). [doi:10.1117/1.3652896].

12 P. Tang, M. Kirby, N. Le, et al., “Polarization sensitive optical coherence tomography with single input for imaging depth-resolved collagen organizations,” Light Sci Appl. 10(1), 237 (2021). [doi:10.1038/s41377-021-00679-3].

13 T. Cannon, N. Uribe-Patarroyo, M. Villiger, et al., “Measuring collagen injury depth for burn severity determination using polarization sensitive optical coherence tomography,” Sci Rep. 12(1), 10479 (2022). [doi:10.1038/s41598-022-14326-3].

14 L. Wang, H. Geng, Y. Liu, et al., “Hot and cold tumors: Immunological features and the therapeutic strategies,” MedComm. 4(5), e343 (2020). [doi:10.1002/mco2.343].

15 F. Kai, A. Drain, and V. Weaver, “The extracellular matrix modulates the metastatic journey,” Dev Cell. 49(3), 332–346 (2019). [doi:10.1016/j.devcel.2019.03.026].

16 M. Chaplain, Multiscale Modelling of Cancer: Micro-, Meso- and Macro-scales of Growth and Spread, 149–168. Springer International Publishing, Cham (2020).

17 Y. Wu and L. Xie, “Ai-driven multi-omics integration for multi-scale predictive modeling of genotype-environment-phenotype relationships,” Computational and Structural Biotechnology Journal. 27, 265–277 (2025). 10.1016/j.csbj.2024.12.030.

18 F. Wang and A. Preininger, “Ai in health: State of the art, challenges, and future directions,” Yearb Med Inform. 28, 16–26 (2025). [doi:10.1055/s-0039-1677908].

19 H. Chan, R. Samala, L. Hadjiiski, et al., “Deep learning in medical image analysis,” Adv Exp Med Biol. 1213, 3–21 (2020).

20 P. Prasetyanti and J. Medema, “Intra-tumor heterogeneity from a cancer stem cell perspective,” Mol Cancer. 16(1), 41 (2017). [doi:10.1186/s12943-017-0600-4].

21 M. Haffner, W. Zwart, M. Roudier, et al., “Genomic and phenotypic heterogeneity in prostate cancer,” Nat Rev Urol. 18(2), 79–92 (2021). 10.1038/s41585-020-00400-w.

22 M. Chen, S. Copley, P. Viola, et al., “Radiomics and artificial intelligence for precision medicine in lung cancer treatment,” Semin Cancer Biol. 93, 97113 (2025). [doi:10.1016/j.semcancer.2023.05.004].

23 A. Schroeder, O. Mueller, S. Stocker, et al., “The rin: an rna integrity number for assigning integrity values to rna measurements,” BMC Mol Biol. 7, 3 (2006). [doi:10.1186/1471-2199-7-3].

24 T. Matsubara, J. Soh, M. Morita, et al., “Dv200 index for assessing rna integrity in next-generation sequencing,” Biomed Res Int. 2020, 9349132 (2020). [doi:10.1155/2020/9349132].

25 T. Knapp, S. Duan, J. Merchant, et al., “Quantitative characterization of duodenal gastrinoma autofluorescence using multi-photon microscopy,” Lasers Surg Med. 55(2), 208–225 (2023). [doi:10.1002/lsm.23619].

26 T. Knapp, N. Lima, N. Daigle, et al., “Combined flat-field and frequency filter approach to correcting artifacts of multichannel two-photon microscopy,” J Biomed Opt. 29(1), 016007 (2025). [doi:10.1117/1.JBO.29.1.016007].

27 A. Cardona, S. Saalfeld, J. Schindelin, et al., “Trakem2 software for neural circuit reconstruction.,” PLoS One. 7, 38011 (2012). [doi:10.1371/journal.pone.0038011].

28 S. Saalfeld, A. Cardona, V. Hartenstein, et al., “As-rigid-as-possible mosaicking and serial section registration of large sstem datasets,” Bioinformatics. 26, i57–i63 (2010). [doi:10.1093/bioinformatics/btq219].

29 E. Zormpas, R. Queen, A. Comber, et al., “Mapping the transcriptome: Realizing the full potential of spatial data analysis,” Cell. 186, 5677 – 5689 (2023). 10.1016/j.cell.2023.11.003.

30 R. M. Haralick, K. Shanmugam, and I. Dinstein, “Textural features for image classification,” *IEEE Transactions on Systems*, Man, and Cybernetics SMC-3(6), 610–621 (1973).

31 K. Simonyan and A. Zisserman, “Very deep convolutional networks for large-scale image recognition,” arXiv: 1409.1556v6 (2015). 10.48550/arXiv.1409.1556.

32 A. Weisbrod, L. Zhang, M. Jain, et al., “Altered pten, atrx, chga, chgb, and tp53 expression are associated with aggressive vhl-associated pancreatic neuroendocrine tumors,” Horm Cancer. 4, 165–175 (2013). [doi:10.1007/s12672-013-0134-1].

33 S. Duan, T. Sawyer, R. Sontz, et al., “Gfap-directed inactivation of men1 exploits glial cell plasticity in favor of neuroendocrine reprogramming,” Cell Mol Gastroenterol Hepatol. 14(5), 1025–1051 (2022).

34 C. Shelton, S. Solomon, J. LaRusch, et al., “Prss1-related hereditary pancreatitis,” GeneReviews (2012).

35 J. Hofland, M. Falconi, E. Christ, et al., “European neuroendocrine tumor society 2023 guidance paper for functioning pancreatic neuroendocrine tumour syndromes,” J Neuroendocrinol. 35, e13318 (2023). [doi:10.1111/jne.13318].

36 R. Maréchal, J.-B. Bachet, A. Calomme, et al., “Sonic hedgehog and gli1 expression predict outcome in resected pancreatic adenocarcinoma,” Clin Cancer Res. 21, 1215–1224 (2015). 10.1158/1078-0432.CCR-14-0667.

37 K. Taki, D. Hashimoto, S. Nakagawa, et al., “Significance of lymph node metastasis in pancreatic neuroendocrine tumor,” Surg Today. 47, 1104–1110 (2017). [doi:10.1007/s00595-017-1485-y].

38 T. Cannon, J. Lagarto, B. Dyer, et al., “Characterization of nadh fluorescence properties under one-photon excitation with respect to temperature, ph, and binding to lactate dehydrogenase,” OSA Contin. 4, 1610–1625 (2021). [doi:10.1364/OSAC.423082].

39 A. Sánchez-Hernández, C. Polleys, and I. Georgakoudi, “Formalin fixation and paraffin embedding interfere with preservation of optical metabolic assessments based on endogenous nad(p)h and fad two photon excited fluorescence,” Preprint. bioRxiv., 545363 (2023). [doi:10.1101/2023.06.16.545363].

